# Identification of *Arhgef12* and *Prkci* as Genetic Modifiers of Retinal Dysplasia in the *Crb1^rd8^* Mouse Model

**DOI:** 10.1101/2021.09.02.458662

**Authors:** Sonia M. Weatherly, Gayle B. Collin, Jeremy R. Charette, Lisa Stone, Nattaya Damkham, Lillian F. Hyde, James G. Peterson, Wanda Hicks, Gregory W. Carter, Jürgen K. Naggert, Mark P. Krebs, Patsy M. Nishina

## Abstract

Mutations in the apicobasal polarity gene *CRB1* lead to diverse retinal diseases, such as Leber congenital amaurosis, cone-rod dystrophy, retinitis pigmentosa (with and without Coats-like vasculopathy), foveal retinoschisis, macular dystrophy, and pigmented paravenous chorioretinal atrophy. Limited correlation between disease phenotypes and *CRB1* alleles, and evidence that patients sharing the same alleles often present with different disease features, suggest that genetic modifiers contribute to clinical variation. Similarly, the retinal phenotype of mice bearing the *Crb1* retinal degeneration 8 (*rd8*) allele varies with genetic background. Here, we initiated a sensitized chemical mutagenesis screen in B6.Cg-*Crb1^rd8^*/Pjn, a strain with a mild clinical presentation, to identify genetic modifiers that cause a more severe disease phenotype. Two models from this screen, *Tvrm266* and *Tvrm323*, exhibited increased retinal dysplasia. Genetic mapping with high-throughput exome and candidate-gene sequencing identified causative mutations in *Arhgef12* and *Prkci*, respectively. Epistasis analysis of both strains indicated that the increased dysplastic phenotype required homozygosity of the *Crb1^rd8^* allele. Retinal dysplastic lesions in *Tvrm266* mice were smaller and caused less photoreceptor degeneration than those in *Tvrm323* mice, which developed an early, large diffuse lesion phenotype. In both models at one month of age, Müller glia and microglia mislocalization at dysplastic lesions was similar to that in B6.Cg-*Crb1^rd8^*/Pjn mice, while photoreceptor cell mislocalization was more extensive. External limiting membrane disruption was comparable in *Tvrm266* and B6.Cg- *Crb1^rd8^*/Pjn mice but milder in *Tvrm323* mice. Immunohistological analysis of mice at postnatal day 0 indicated a normal distribution of mitotic cells in *Tvrm266* and *Tvrm323* mice, suggesting normal early development. Aberrant electroretinography responses were observed in both models but functional decline was significant only in *Tvrm323* mice. These results identify *Arhgef12* and *Prkci* as modifier genes that differentially shape *Crb1-*associated retinal disease, which may be relevant to understanding clinical variability and underlying disease mechanisms.

## Introduction

Inherited retinal dystrophies associated with variants of the apicobasal polarity gene *CRB1* exhibit a perplexing diversity of disease phenotypes (reviewed in [1–6]), including Leber congenital amaurosis (LCA8, MIM 613835), early-onset rod-cone dystrophy, juvenile- or adult-onset retinitis pigmentosa (RP) with or without paraarteriolar preservation of the retinal pigment epithelium (RP12, MIM 600105), cone-rod dystrophy, RP with Coats-like exudative vasculopathy (retinal telangiectasia), hypermetropia, keratoconus, foveal retinoschisis and cystic or retinoschisis-like maculopathy and macular dystrophy, and pigmented paravenous chorioretinal atrophy (PPCRA, MIM 172870). This variability in disease onset, progression, severity, topography, and specific pathological features makes it difficult to advise patients about therapeutic options and key quality of life issues, and to identify suitable patients for participation in prospective clinical trials [7]. Identifying the genetic and/or environmental factors responsible for clinical variability may therefore refine the prognosis of *CRB1-*associated diseases and promote the development of therapeutic approaches.

Clinical variability may reflect the many biological processes in which CRB1 participates. *CRB1* encodes a mammalian member [8] of a family of transmembrane proteins related to *Drosophila* Crumbs (Crb) [9], which are central to conserved CRB complexes that govern epithelial apicobasal polarity, cell-cell adhesion, apical segregation of proteins and lipids, cellular size and shape determination, intercellular signaling, cell division, and tissue morphogenesis [4], [10–14]. In the *Drosophila* eye, Crb localizes to the stalk subdomain of the photoreceptor cell apical membrane, where it mediates the assembly of adherens junctions and the apical segregation of cellular components [13], [15–17]. Homozygous mutations of *crb* disrupt photoreceptor cell morphogenesis and cause photoreceptor cell apoptosis and retinal degeneration [16], [18]. In the mouse and human retina, CRB1 is most prominently localized to the subapical region of the neuroepithelium, specifically in photoreceptor inner segments above adherens and tight junctions, and in Müller cell apical processes [8], [16], [19], [20]. CRB1 disruption in mice perturbs cell-cell adhesion at the external limiting membrane (ELM), as indicated by the focal loss of adherens junctions [19], [21]. CRB1 disruption is also associated with outer retinal folds and pseudorosettes (retinal dysplasia) that correspond to light spots observed by fundus examination, and with photoreceptor degeneration in dysplastic regions [19–23]. Thus, CRB1 engages in multiple activities affecting cell and tissue integrity, raising the possibility that the variability of *CRB1/Crb1-*associated retinal disease arises from the differential disruption of these activities.

One hypothesis to explain disease variability is that *CRB1* variants have allele-specific effects. Nearly 300 pathogenic or likely pathogenic *CRB1* alleles have been identified (https://databases.lovd.nl/shared/genes/CRB1). Alleles may differ in their effect on protein levels, stability, or function, and variants affecting specific protein domains might be associated with distinct *CRB1-*disease subtypes. Although most efforts to establish genotype-phenotype correlations have met with limited success to date [1–6], some recent clinical studies indicate potential correlations [7], [24], [25]. The correlation of disease-causing missense substitutions in the extracellular domains of *Drosophila* Crb with variable cellular and retinal degeneration phenotypes [26] and the recent identification of a novel photoreceptor-specific isoform, CRB1- B, in human and mouse retinas [21], may provide new perspectives for understanding these allele-specific effects.

It has also been suggested that variation in modifier genes may contribute to *CRB1-*associated disease variability. For example, mutations in genes encoding proteins that may directly or indirectly interact with CRB1 might mediate different pathogenic effects. In support of this hypothesis, distinct and variable disease phenotypes have been observed among individuals with identical *CRB1* alleles, suggesting the existence of genetic modifiers [2–4], [6], [7], [27–29].

Additionally, in about 30% of affected individuals only a single *CRB1* variant allele has been detected suggesting that other genetic variants, possibly including modifier loci, may contribute to the disease [2]. However, apart from a report that an allele of the LCA and RP gene, *AIPL1,* acted as a potential modifier of *CRB1* retinal pathology [30], modifiers in the human population have not been identified. Identification and validation of modifier genes is challenging given the limited number of affected individuals for rare diseases, such as those associated with *CRB1* variants, the genetic heterogeneity of the human population, and the possible effects of undefined environmental factors on disease phenotype.

The successful use of mouse models to identify genetic modifiers of human diseases [31], [32] extends to models of eye diseases [33], including *CRB1-*associated retinal dystrophy. Early evidence for genetic modifiers in a *CRB1-*disease model was obtained from studies of STOCK *Crb1^rd8^*. This genetically mixed inbred strain exhibited extensive dysplasia in the inferior retina observed as light spots by indirect ophthalmoscopy and fundus imaging, which correspond to outer retinal pseudorosettes and folds [19], [34]. Backcrossing STOCK *Crb1^rd8^* with the wild-type strains CAST/EiJ or C57BL/6J (B6) suppressed the dysplastic phenotype [19], [34], indicating an effect of genetic background. Further evidence for modifiers of the phenotype was obtained from breeding studies of inbred *Crb1^rd8^* strains [35]. Variability of the dysplastic phenotype was also noted among mice derived from the C57BL/6N (B6N) substrain, which is homozygous for the *Crb1^rd8^* allele [36], and in strains from The Jackson Laboratory (JAX) collection [37]. *Cx3cr1, Mthfr,* and *Cygb* were identified as modifiers of *Crb1^rd8^* retinal dysplasia [38–40], *Jak3* as an enhancer of a *Crb1^rd8^*-dependent neovascular phenotype [41], and *Crb2* as a modifier of retinal dysplasia due to a *Crb1* knock-out allele [42], [43]. Further, the choroidal neovascularization phenotype of an *Nfe2l2* knock-out strain [44] was more severe in the presence of homozygous *Crb1^rd8^* alleles [45], indicating gene interaction. These studies reveal substantial variability in the retinal phenotype of *Crb1* mutant mice depending on genetic background and demonstrate that modifier genes can be identified. However, the modifiers identified so far do not appear to participate in shared pathways, so identification of additional modifier genes is needed to reveal the cellular and molecular networks that account for all of the observed *CRB1-*disease variability.

Here, as part of the Translational Vision Research Models (TVRM) program [46–48], we identified genetic modifiers that cause a more severe *Crb1^rd8^* dysplastic retinal phenotype using a sensitized *N*-ethyl-*N*-nitrosourea (ENU) mutagenesis screen of B6.Cg-*Crb1^rd8^*/Pjn (hereafter B6 *rd8*) mice. These mice are congenic on the B6 background, homozygous for the *Crb1^rd8^* allele, and exhibit a near-normal fundus appearance [19]. Two mutant modifier lines, *Tvrm266* and *Tvrm323*, were found, respectively, to carry mutations in *Arhgef12,* which encodes guanine nucleotide exchange factor 12, a modulator of Rho GTPase activity, and *Prkci*, which encodes a protein homologous to *Drosophila* atypical protein kinase C, a component of the Baz-aPKC-Par- 6 polarity complex that engages with Crb [10], [49]. This work extends the network of apicobasal polarity components that influence *CRB1-*associated retinal disease and underscores its complexity and interconnectivity with multiple molecules and pathways.

## Results

### Sensitized Chemical Mutagenesis Screen of *Crb1^rd8^* Mice

Identification of variants contributing to the severe dysplastic phenotype in STOCK *Crb1^rd8^* mice by classic genetical means of recombinant mapping proved to be inconclusive, likely due to multigenic effects and gene interactions. Therefore, to identify genetic modifiers of *Crb1^rd8^* dysplasia, two sensitized mutagenesis screens were considered initially: mutagenesis of STOCK *Crb1^rd8^* mice and screen for decreased severity of the fundus spotting phenotype, or mutagenesis of B6 *rd8*, an incipient congenic strain with a mild phenotype [19], and screen for increased severity of the phenotype. Mutagenesis of STOCK *Crb1^rd8^* males failed to yield pups in repeated attempts with varying concentrations and timing of ENU injections. Thus, mutagenized B6 *rd8* males were bred according to an established strategy to detect both dominant and recessive mutations [46], and the resulting G_3_ population screened for an increased spotting phenotype (Fig. 1).

**Fig. 1.**
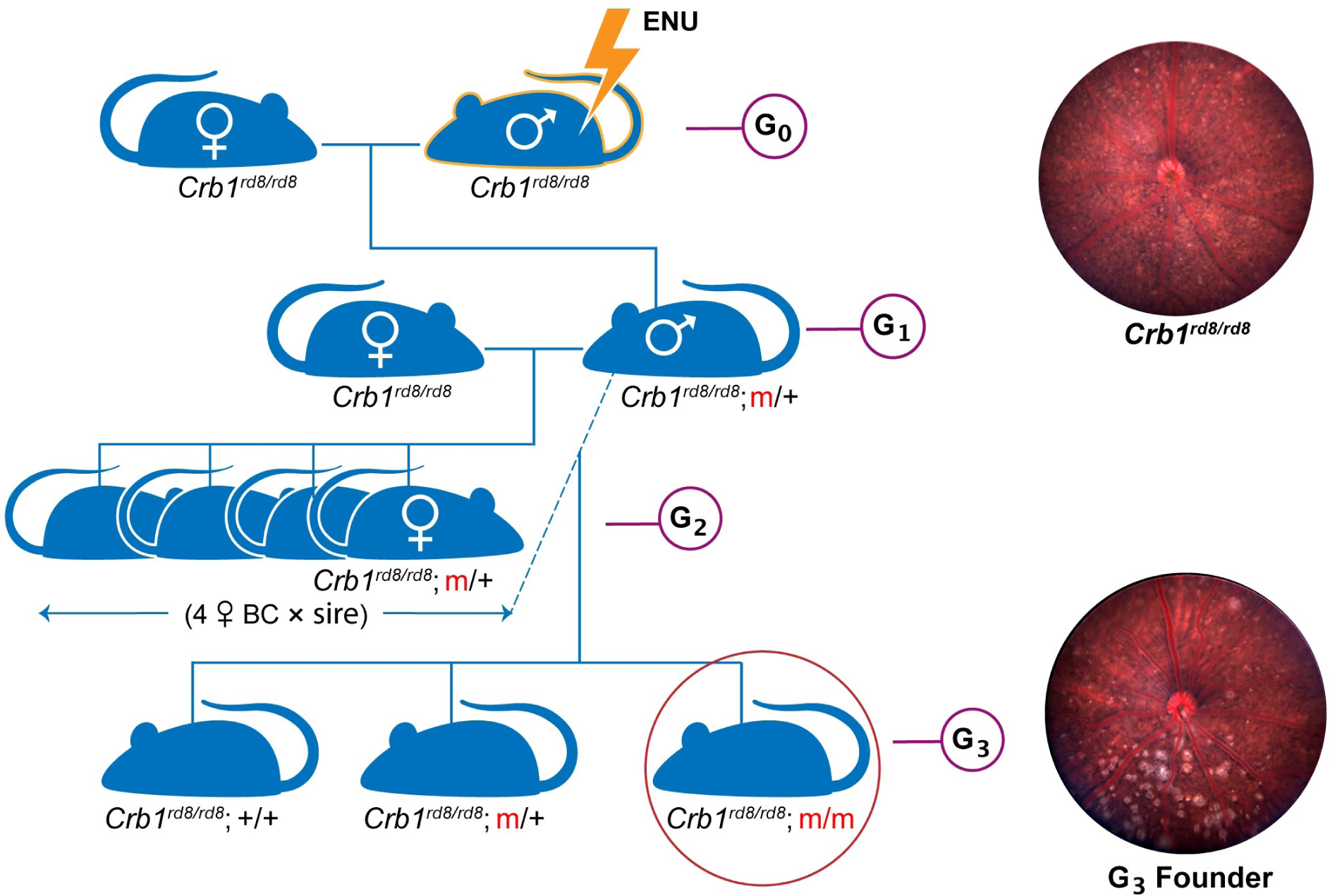
Breeding strategy for chemical mutagenesis. B6 *rd8* G_0_ males were treated with N-ethyl-N-nitrosourea (ENU). After about three months when G_0_ males regained fertility, they were mated to B6 *rd8* females to produce G_1_ progeny. G_1_ males were mated to produce G_2_ progeny, of which four females were backcrossed with the same G_1_ male to produce G_3_ progeny. G_3_ mice were screened by indirect ophthalmoscopy for evidence of increased retinal dysplasia. G_3_ founders were developed into TVRM models by testing for heritability and further backcrossing to remove unlinked mutagenized genes. The increased spotting phenotype typical of retinal dysplasia is evident by comparing the fundus images of a B6 *rd8* mouse (G_0_) and a G_3_ founder at 12 weeks of age.

### Identification of mutations

Two mutant lines, *Tvrm266* and *Tvrm323*, exhibited a heritable, bilateral increase in fundus spots compared to the parental B6 *rd8* mice (Fig. 2A) and were backcrossed for a minimum of five generations to unmutagenized B6 *rd8* to remove unlinked mutations prior to characterization. A high fraction of affected progeny during initial intercrosses to expand the colony suggested that the phenotype in both strains might be semi-dominant. The modifier loci were mapped by crossing these strains with B6N mice, which are homozygous for the *Crb1^rd8^* allele [36] and have diverged sufficiently from B6 to allow for chromosomal mapping of the modifying loci. For *Tvrm266* mice, high-throughput whole-exome sequencing of affected individuals was first performed to identify likely mutation candidates. For *Tvrm323* mice, a genome-wide recombinational mapping screen (S1 Fig, S1 Data) of 103 backcross progeny ((*Tvrm323* × B6N) F_1_ × B6N) revealed that the interval yielding the highest frequency of affected heterozygotes (72%) was a 15.3 Mbp region of Chromosome 3 (Chr 3; 03-021837059 to 03-037129353).

**Fig. 2.**
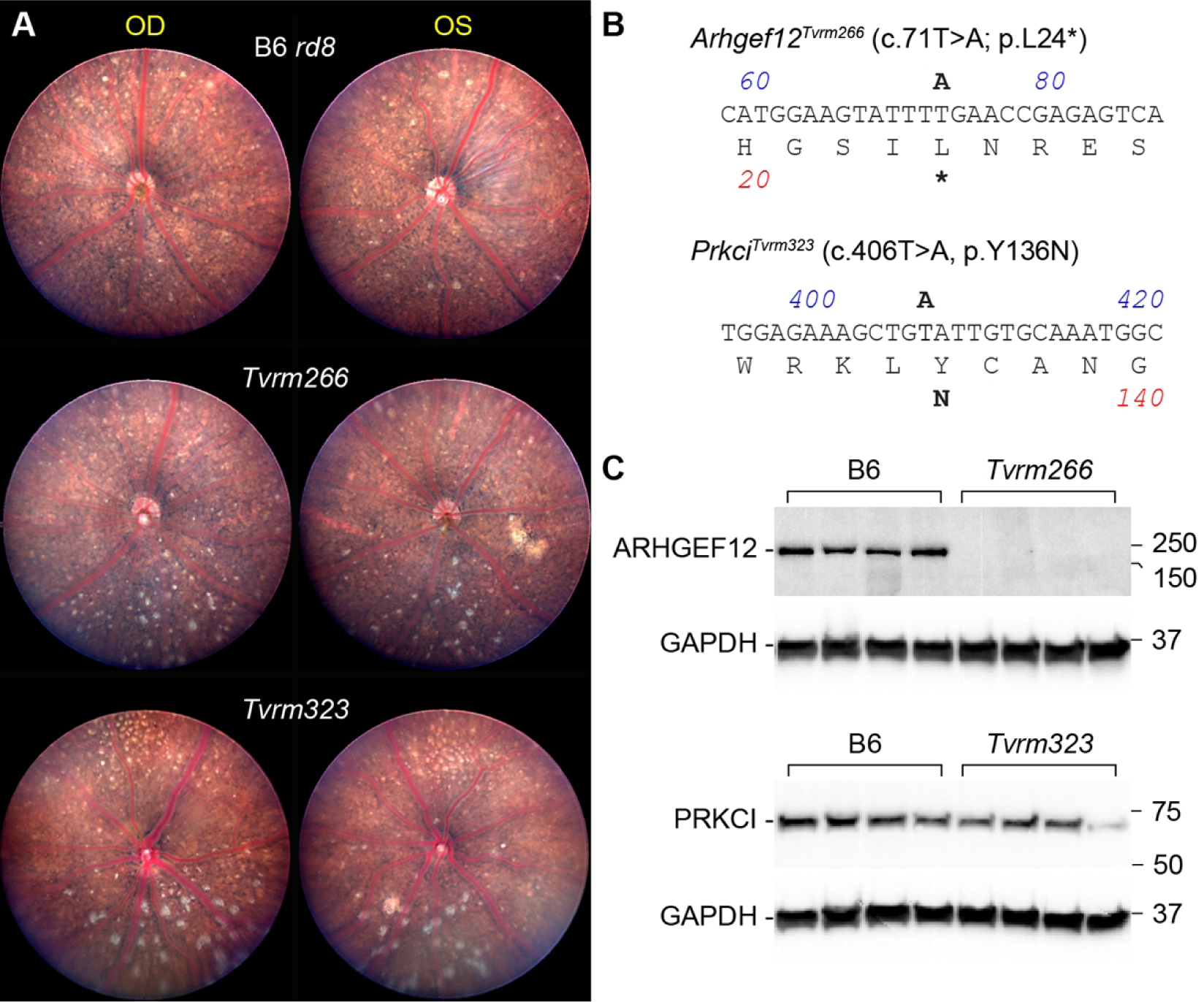
Identification of mutations and protein expression in modifier strains. A. Fundus photodocumentation of B6 *rd8* and *Tvrm266* mice at 10 weeks of age, and *Tvrm323* mice at 3 months of age. Right (OD) and left (OS) eyes of a single mouse are shown. The lower portion of each image corresponds to the inferior retina based on the position of the mouse head relative to the camera during imaging. B. Sequence and amino acid translation of the mutant alleles. The DNA sequence corresponds to a portion of the forward strand of each gene, numbered as in the mRNA. Mutations are indicated in *bold* above the sequence. The predicted amino acid sequence is given beneath the DNA sequence using the single-letter code and the predicted translational effect of the mutation is indicated in *bold* below. C. Enhanced chemiluminescence western analysis. Posterior eyecup lysates from B6 control and modifier strains (n = 4) were electrophoresed and transferred to nitrocellulose. *Tvrm266* blots were probed with ARHGEF12 antibody and *Tvrm323* blots with PRKCI antibody. After imaging, blots were probed with GAPDH antibody and reimaged to control for protein recovery and loading. Molecular weights of protein standards are indicated in kDa.

Putative causative mutations were identified in *Arhgef12*, by candidate gene sequencing from the whole-exome sequences of *Tvrm266* mice, and in *Prkci,* by candidate gene sequencing within the confidence interval identified by mapping *Tvrm323* mice (Fig. 2B, S2 Fig). Co-segregation of disease phenotype with the mutations identified was statistically significant by endpoint genotyping analysis of 16 progeny from a *Tvrm266* mapping cross (*Tvrm266* × B6N) F_1_ × B6N; *p* = 0.010, Fisher’s Exact Test) and by sequencing 12 progeny from the *Tvrm323* mapping cross (*p* = 5.4 × 10^-5^, Fisher’s Exact Test), indicating that these mutations are causative (S2 Fig).

To assess the effect of the mutations on expression of the encoded proteins, western analysis was performed (Fig. 2C). Analysis of control B6 samples revealed a single band consistent with the predicted molecular weights of ARHGEF12 (171 kDa predicted for isoform X1, XP_017169054.1) and PRKCI (65 kDa predicted for isoform X1, XP_006535474). Quantitation of the western blots (S3 Fig, S1 Data) indicated that the ARHGEF12 level in *Tvrm266* eyes was no more than 0.037 of that in B6 eyecups (Student’s t-test, *p* = 0.0015); this value is likely to be an overestimate arising from non-specific staining of other eyecup proteins in the blot region analyzed. Little to no expression is expected due to the premature stop codon caused by the mutation. PRKCI levels in *Tvrm323* mice were 0.57 of those in B6 mice but this effect was not statistically significant (Student’s t-test, *p* = 0.052). Comparison of B6 *rd8*, *Tvrm266*, and *Tvrm323* to B6 control eyecups by qRT-PCR (S1 Data) indicated statistically significant differences in *Arhgef12* mRNA levels among strains (one-way ANOVA of Δ*C_t_* values, F[3, 20] = 6.869, *p* = 0.0023). Dunnet’s *post-hoc* test revealed a significant decrease in *Arhgef12* mRNA levels (0.74 of B6, *p* = 0.0018) but not in other strains. This decrease may reflect nonsense- mediated decay of a subset of *Arhgef12* transcripts in *Tvrm266* mice. Parallel analysis of *Prkci* mRNA did not indicate significant *post hoc* differences among the strains compared to B6 controls (S1 Data). Taken together, these results establish novel alleles of two genes as candidate modifiers of the *Crb1^rd8^* fundus phenotype. The full designation for these models is B6.Cg-*Crb1^rd8^ Arhgef12^Tvrm266^*/Pjn and B6.Cg-*Crb1^rd8^ Prkci^Tvrm323^*/Pjn; for simplicity, we retain the original strain designations *Tvrm266* and *Tvrm323*, respectively, for the remainder of the paper, and unless otherwise indicated the genotype is homozygous for both *Crb1^rd8^* and the modifier.

### Epistasis of *Crb1^rd8^* modifier genes

We postulated that the increased retinal spotting phenotypes of *Tvrm266* and *Tvrm323* may be due to the new mutations acting alone or through an interaction with *Crb1^rd8^*. To distinguish these possibilities, we tested for epistasis, classically defined as the masking of a phenotype associated with a variant at one locus by a variant at a second [50]. To examine epistatic interactions, each modifier strain was outcrossed to B6 mice, which are wild-type for *Crb1^rd8^*, and subsequently intercrossed. In the case of *Tvrm266*, F_2_ mice progeny were genotyped and intercrossed to produce F_3_ mice of select genotypes for additional analysis. Increased fundus spotting was observed in heterozygous or homozygous *Arhgef12^Tvrm266^* or *Prkci^Tvrm323^* mice in the presence of the homozygous, but not heterozygous, *Crb1^rd8^* allele (Fig. 3A). The fundus phenotype was more severe in the presence of two alleles (homozygous) than with one allele (heterozygous) of either *Arhgef12^Tvrm266^* or *Prkci^Tvrm323^*. These results were extended by indirect ophthalmoscopy to all nine genotypes expected from the epistasis intercross (Fig. 3B and C).

**Fig. 3.**
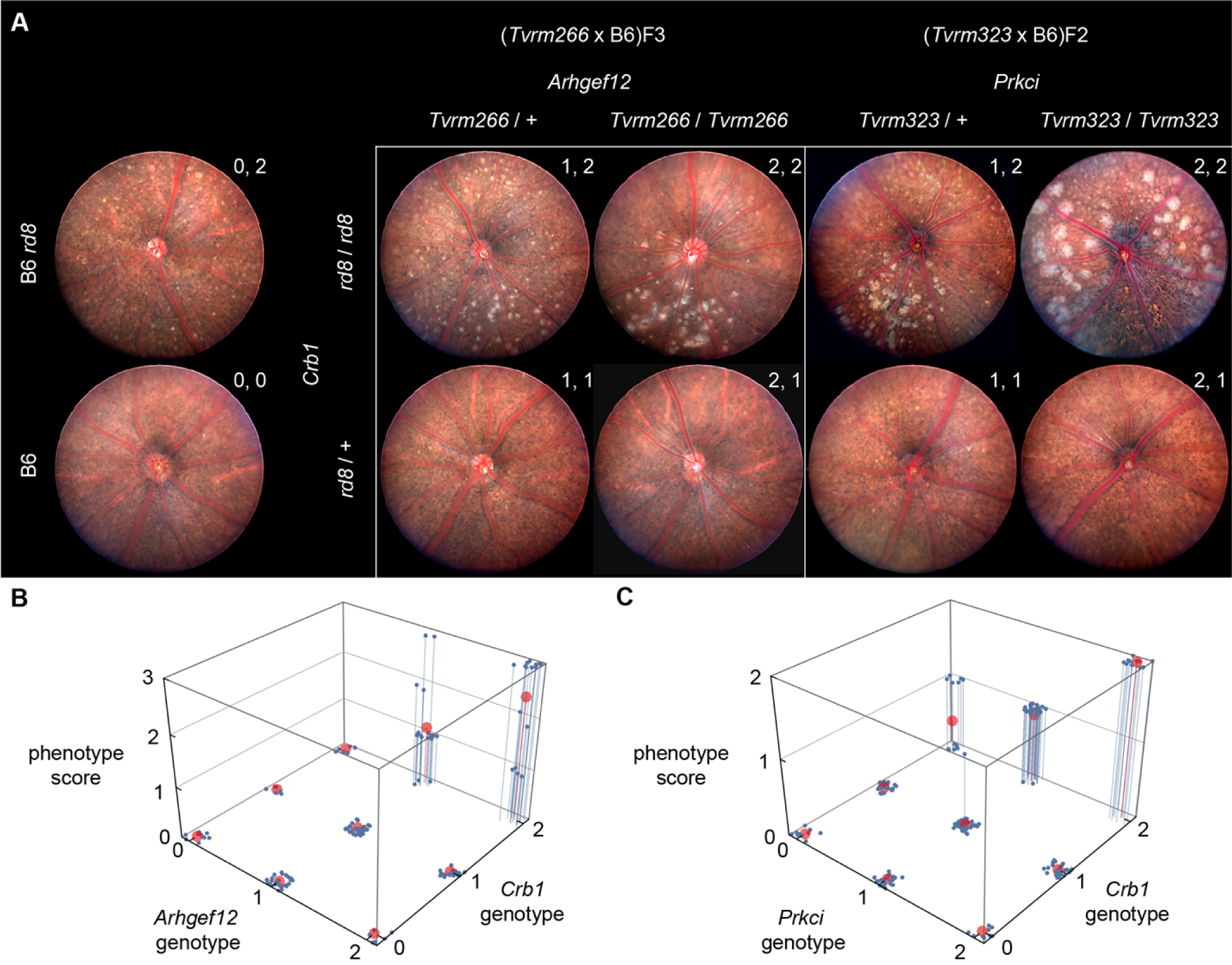
A. Fundus photographs of B6, B6 *rd8*, and F_3_ progeny from the *Tvrm266* epistasis cross at 10 weeks of age or F_2_ progeny from the *Tvrm323* epistasis cross at 2.5–3.7 months of age. Genotypes are indicated by m/+ (heterozygous) or m/m (homozygous), where m is the mutant allele symbol and, for comparison with panels B and C, by the values a, b at the upper right of each fundus image, which indicate the number of mutant alleles at the modifier and *Crb1* loci, respectively (0, wild-type; 1, heterozygous; 2, homozygous). At least 3 females and 3 males of each genotype were imaged bilaterally. B. 3D plot of the indirect ophthalmoscopy phenotypes of F_2_ and F_3_ progeny from the *Tvrm266* epistasis cross (9–15 weeks of age) against the nine genotypes expected (n = 6–21 for each genotype). C. Corresponding plot of F_2_ phenotypes from the *Tvrm323* epistasis cross (2.5–3.7 months of age; n = 10–48 for each genotype). Indirect ophthalmoscopy annotations were converted to ordinal phenotype scores. Scores for individual mice of each genotype examined are shown in *blue*, with fit statistical models represented by *red circles*. Genotypes values are indicated as in panel A.

Phenotypes of heterozygous *Crb1^rd8^* mice were similar to those of B6 mice, which rarely exhibited spots (Fig. 3A). Interaction modeling of indirect ophthalmoscopy data (S1 Data) from epistasis crosses of both mutants indicated a significantly improved fit using a model that accounted for gene-gene interaction (Fig. 3B and C) compared to one that was purely additive (adjusted *R^2^* value for interacting vs. additive models: *Tvrm266* cross, 0.78 vs. 0.62; *Tvrm323* cross, 0.89 vs. 0.69). An F-ratio test of the F-statistic derived from the two statistical models was significant, indicating superior performance of the interacting model for both mutants (F-statistic and *p* values: *Tvrm266* cross, 26.1 and *p* = 6.3 × 10^-16^; *Tvrm323* cross, 97.9 and *p* = 5.4 × 10^-46^). Both analyses revealed a significant effect of the heterozygous allele on the phenotype (*Tvrm266* cross, *p* = 1.9 × 10^-5^; *Tvrm323* cross*, p* = 1.6 × 10^-8^), which was smaller than that of the homozygous allele, confirming the semi-dominant mode of inheritance.

Taken together, these results reveal a semi-dominant epistatic interaction between *Arhgef12* or *Prkci* mutant alleles and the *Crb1^rd8^* mutation leading to increased fundus spotting.

### Pathological features of modifier strains

Having established that the *Arhgef12* and *Prkci* alleles modify the *Crb1^rd8^* phenotype, we sought to gain insights into disease pathogenesis by studying the mutant strains. Analysis was performed on homozygous *Tvrm266* and *Tvrm323* mice, as the phenotype could be observed at one month of age. To determine whether increased fundus spotting was correlated with dysplastic lesions and other pathological changes, eyes collected from B6, B6 *rd8*, *Tvrm266*, and *Tvrm323* mice were examined histologically (Fig. 4). We defined dysplastic lesions as retinal folds or perturbations resulting in the displacement of photoreceptor nuclei by more than three nuclear diameters from either boundary of the outer nuclear layer (ONL). B6 *rd8* retinas (Fig. 4B) were morphologically similar to B6 retinas (Fig. 4A) except for rare dysplastic lesions in the inferior retina (Fig. 4B, *asterisk*; also compare the higher magnification images in Fig. 4E and F). By contrast, retinal morphology was disrupted more extensively in the modifier strains, as reported previously in STOCK *Crb1^rd8^* mice [19], [23], [34] and other *Crb1* mutant models [20]. Lesions were also evident in the superior retina of *Tvrm266* and *Tvrm323* near the optic nerve head. Dysplastic lesions were larger than those in B6 *rd8* mice (compare Fig. 4G, H with Fig. 4F) and differed between the two modifier strains. In *Tvrm266* mice, the largest lesions (Fig. 4C, *asterisks*; Fig. 4G) extended toward the inner nuclear layer (INL) and resembled the pseudorosettes described in STOCK *Crb1^rd8^* mice and other models [19], [20], [23], [34]. By contrast, *Tvrm323* lesions were characterized by displacement of retinal cell layers toward the RPE (Fig. 4D, *asterisks*; Fig. 4H) and often contained tubular structures that reached and perturbed cells of this layer (Fig. 4H, *red arrowhead*). These structures may represent retinal neovessels, a frequently reported attribute of *Crb1* mutant mice and rats [35], [41], [51]. A novel feature discovered in modifier strains at one month of age was the presence of apparent voids in the ONL near dysplastic lesions (Fig. 4G and H, *yellow arrowheads*). One sample with ONL voids was also observed among nine B6 *rd8* mice examined (Fig. 4I).

**Fig. 4.**
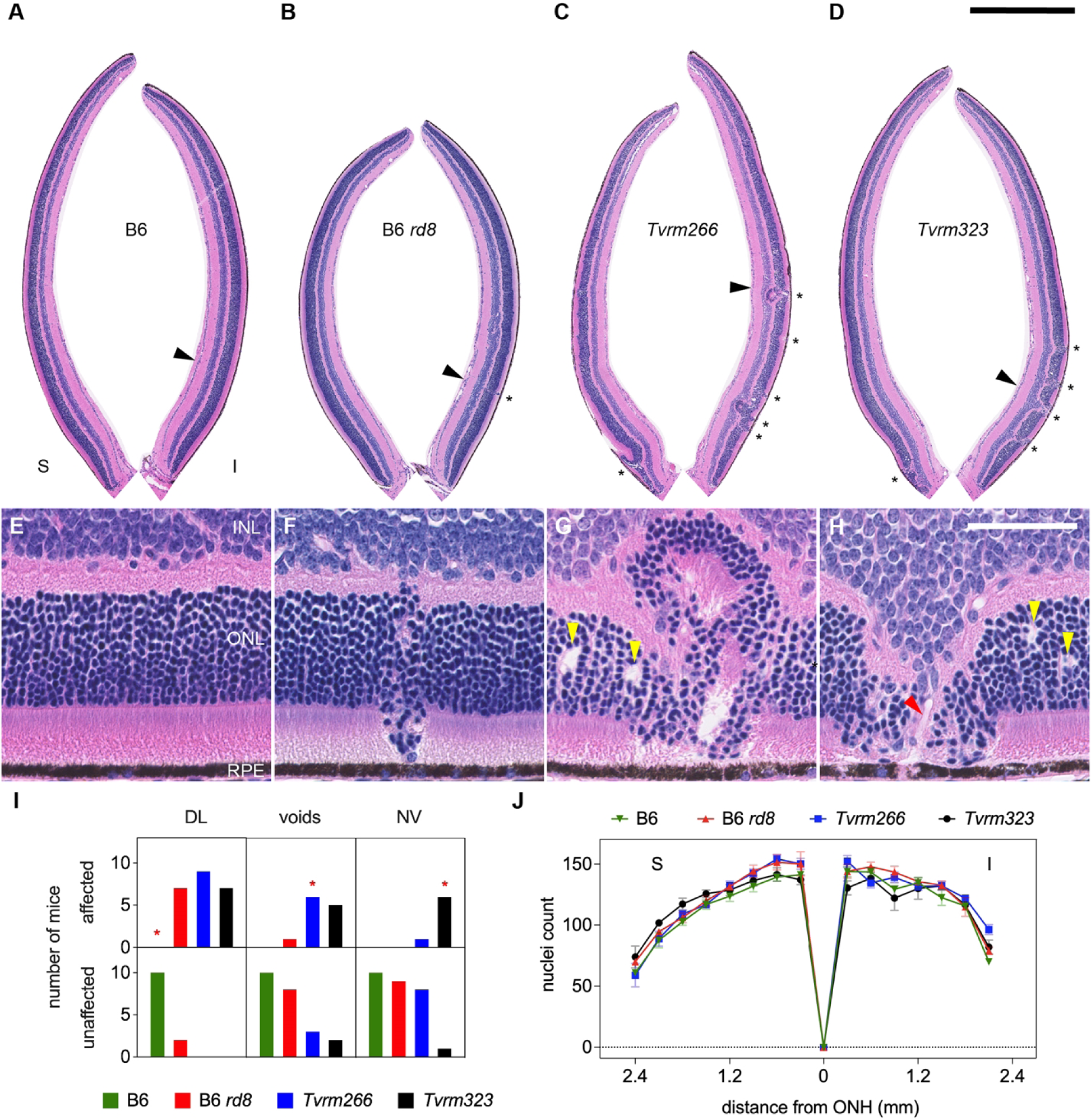
Histological analysis at one month of age. A–D. Superior (S) and inferior (I) retinas cropped from digital images of B6, B6 *rd8*, *Tvrm266* and *Tvrm323* whole eye sections stained with hematoxylin and eosin and imaged with a slide scanner. Dysplastic lesions, *asterisks.* Scale bar, 0.5 mm. Sites from A-D detailed in panels E–H, respectively, are marked by *arrowheads.* E. Normal retinal layers in B6 mice. INL, inner nuclear layer; ONL, outer nuclear layer; RPE, retinal pigment epithelium. F–H. Dysplastic lesions in B6 *rd8, Tvrm266,* and *Tvrm323* mice, respectively. Voids in panels G and H, *yellow arrowheads*. H. Lesion containing possible vascular structure, *red arrowhead.* Scale bar, 50 µm; applies to E–H. I. Incidence of pathological features. Mice showing one or more of the identified features among 17–21 reviewed histological sections were scored as affected. DL, dysplastic lesions; NV, neovessels. J. Photoreceptor nuclei count within a 50 µm length of retina as a function of distance from the ONH. Values represent mean ± standard error of the mean (SEM); n = 5–8.

To assess the statistical significance of these findings, histological sections from each independent sample were reviewed for at least one occurrence of each pathological feature. Compared to B6 *rd8* mice, significant differences in incidence were observed only for dysplastic lesions among B6 mice (Fisher’s exact test, Bonferroni *post hoc* correction, *p* = 0.0021), and for neovascular lesions among *Tvrm323* mice (Fisher’s exact test, Bonferroni *post hoc* correction, *p* = 0.0026; Fig. 4I, *asterisks*; S1 Data). To determine whether these pathological changes were associated with photoreceptor degeneration, photoreceptor nuclei were counted along a fixed length of retina at regular intervals. At one month of age, no significant effect of strain on the number of photoreceptor nuclei was observed among B6 mice or modifier strains compared to B6 *rd8* mice (Fig. 4J, repeated measures mixed-effects model, F (3, 22) = 1.8, *p* = 0.18; S1 Data). These results show that retinal dysplasia and associated pathological changes occur at an early age prior to significant degeneration. The topographical distribution of lesions in the inferior retina and the superior retina near the ONH matched that of fundus spots, indicating that retinal dysplasia accounts for the increased fundus spotting phenotype.

### Age-dependent pathological changes in modifier strains

Most forms of *CRB1* retinal disease are progressive in humans [1–6]. To examine whether progressive changes occurred with age in the modifier strains, histological analysis was repeated at 12 months of age (Fig. 5). Compared to B6 retinas (Fig. 5A, E), B6 *rd8* retinas changed little in aged mice, showing normal morphology with rare dysplastic lesions as observed at one month of age (Fig. 5B, F). By contrast, in addition to a continued presence of dysplastic lesions at 12 months of age (Fig. 5C, D, *asterisks*), retinas in both modifier strains exhibited areas of complete photoreceptor loss, which were more extensive in *Tvrm323* than in *Tvrm266* mice (Fig. 5C, D, *solid lines*). ONL voids were not observed, but neovascular lesions resulting in RPE perturbation were apparent in both modifier strains at this age (Fig. 5G, H, red and *white arrowheads*). The incidence of neovascular lesions was statistically significant in both modifier strains compared to B6 *rd8* (Fig. 5I, S1 Data). Analysis of photoreceptor nuclei counts indicated a statistically significant effect of strain (mixed-effects model, F (3, 36) = 21.42, *p* < 0.0001). In *post hoc* analysis of data from both modifier strains, a statistically significant decrease in nuclei counts was observed near the optic nerve head and extended more peripherally in the inferior than in the superior retina (Fig. 5J, S1 Data). Importantly, degeneration was limited to areas where dysplastic lesions were observed at one month of age (Fig. 4C, D) and the peripheral retina was largely spared, as previously reported in STOCK *Crb1^rd8^* mice [19]. Taken together with the histological findings at one month of age, these results establish that dysplastic lesions, neovascular structures, and photoreceptor cell loss occur and progress to different extents among B6 *rd8*, *Tvrm266,* and *Tvrm323* mice, supporting the hypothesis that modifier genes contribute to *Crb1^rd8^* retinal disease variability.

**Fig. 5.**
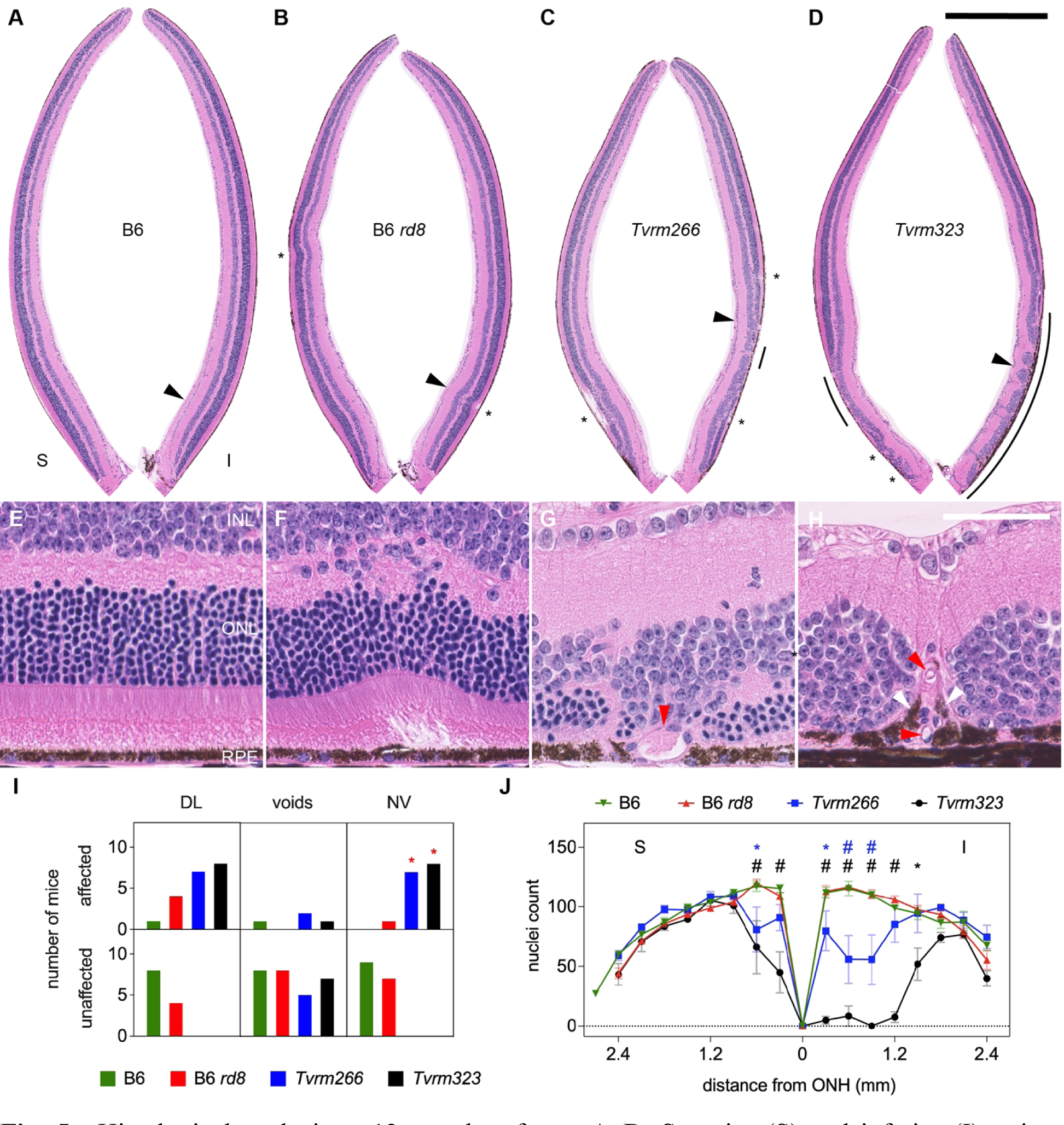
Histological analysis at 12 months of age. A–D. Superior (S) and inferior (I) retinas cropped from digital images of B6, B6 *rd8*, *Tvrm266* and *Tvrm323* whole eye sections stained with hematoxylin and eosin and imaged with a slide scanner. Dysplastic lesions, *asterisks.* Areas devoid of photoreceptor nuclei, *solid lines.* Scale bar, 0.5 mm. Sites from A-D detailed in panels E–H, respectively, are marked by *arrowheads.* E. Normal retinal layers in B6 mice, labeled as in Fig. 4. F. Dysplastic lesion in B6 *rd8.* G, H. Photoreceptor loss at neovascular lesions (*red arrowheads*) in *Tvrm266,* and *Tvrm323* mice, respectively. In G, red blood cells are observed within a large vessel at the RPE. I. Incidence of pathological features among mice. Samples showing one or more of the identified features among 10–21 sections were scored as affected. DL, dysplastic lesions; NV, neovessels. J. Photoreceptor nuclei count within a 50 µm length of retina as a function of distance from the ONH. Values represent mean ± SEM; n = 6–8. Statistical significance colored according to the strain compared to B6 *rd8* in *post hoc* tests. p < 0.01, asterisk; p < 0.0001, hash symbol.

### Cellular disorganization at dysplastic lesions

Retinal dysplasia associated with mutant *Crb1* alleles in mice and rats is characterized by mislocalized photoreceptor cells, shortened photoreceptor outer segments, dysmorphic Müller cells, and the mobilization of immune cells, including microglia and possibly infiltrating macrophages, to the subretinal space [19], [35], [38], [43], [51]. Mislocalized rod photoreceptor cells were readily identified in histological sections, as the nuclei of these cells in adult mice are compact and stain intensely with hematoxylin (Fig. 4). To assess rod outer segment integrity and to test if other retinal cells were abnormally distributed, immunofluorescence analysis of retinal sections was performed using antibody markers for photoreceptor outer segments, Müller glia, and microglia (Fig. 6). Rod and cone outer segments were correctly localized to the outer retina in B6 mice (Fig. 6A) and in intact retinal regions of B6 *rd8, Tvrm266,* and *Tvrm323* mice (Fig. 6B–D). By contrast, outer segments were absent from the RPE interface at dysplastic lesions in all three mutant strains. Small foci of rhodopsin (RHO) and M opsin (OPN1MW) staining were observed throughout the dysplastic areas, consistent with mislocalization of photoreceptor cells. In some instances, intact cone outer segments were observed in the interior of pseudorosettes (Fig. 6C, *arrowheads*), suggesting infolding of the intact outer retina. In B6 mice, Müller glia were observed with soma positioned at the center of the INL and retina-spanning processes extending to endfeet at the internal limiting membrane (ILM) and ELM (Fig. 6E, I). Similar morphology was observed in unaffected regions of B6 *rd8, Tvrm266,* and *Tvrm323* retinas (Fig. 6F–H,), but at dysplastic lesions Müller cell bodies were occasionally displaced toward the ONL (Fig. 6F–H, J–L *arrowheads*). Microglia were found in their normal locations on the outer surface of the INL in B6 mice (Fig. 6M) and in unaffected regions of the three *Crb1^rd8^* strains (Fig. 6N–P). By contrast, at dysplastic lesions of these strains, microglial cells accumulated in the subretinal space or appeared to be moving toward this region (Fig. 6N–P). As in other *Crb1* mouse and rat models [19], [35], [38], [43], [51], these results emphasize an enhanced focal pathology in *Tvrm266* and *Tvrm323* mice that is limited to dysplastic lesions, in which photoreceptor cells are displaced toward the INL from their normal locations and Müller cells and microglia are mobilized toward the outer retina and RPE.

**Fig. 6.**
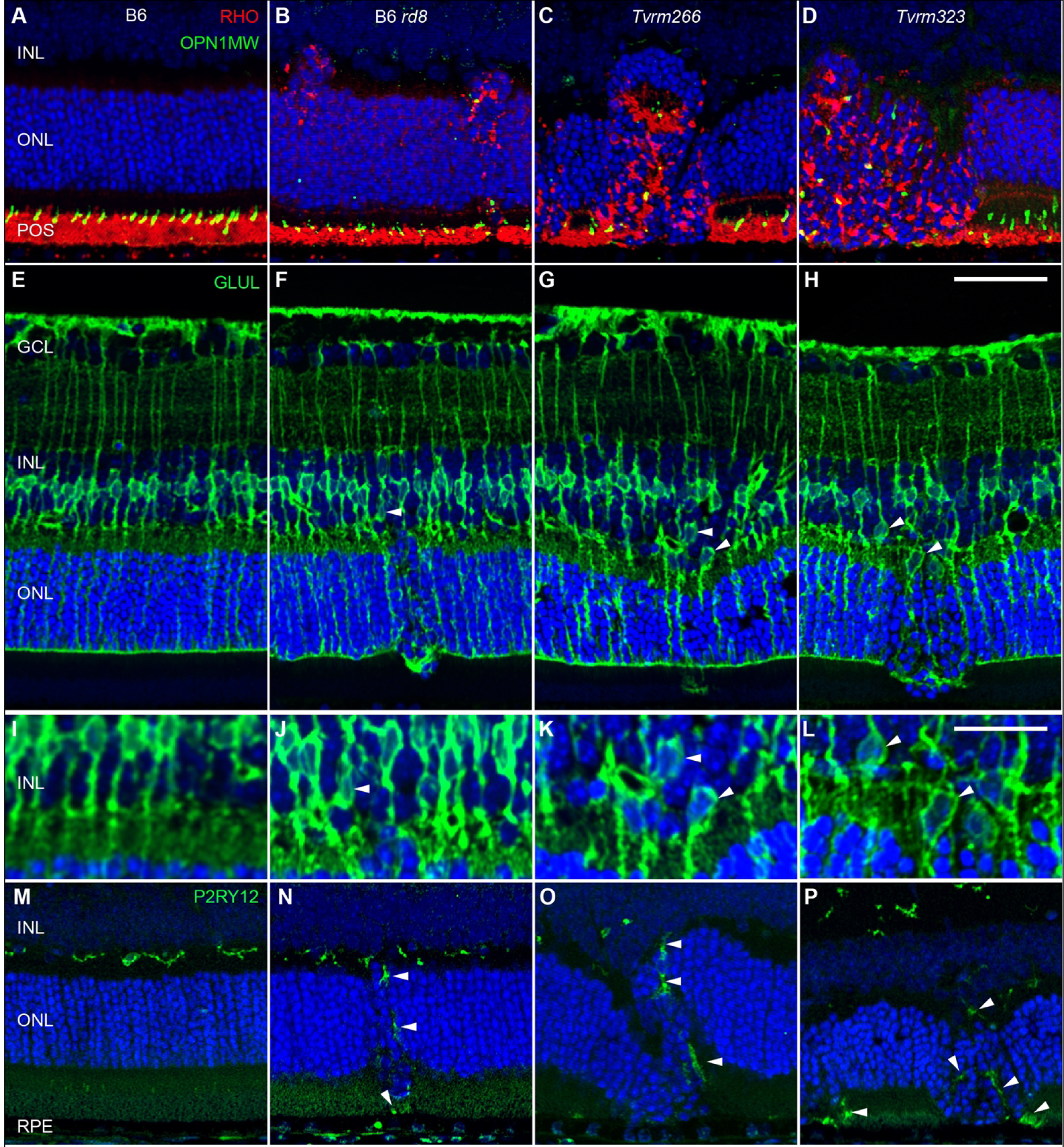
Ectopic retinal cells in dysplastic lesions at one month of age. Retinal sections of 7–9 B6 (A, E, I, M), B6 *rd8* (B, F, J, N), *Tvrm266* (C, G, K, O), and *Tvrm323* (D, H, L, P) mice were examined for each panel of markers studied. A–D. Staining with rhodopsin (RHO, *red*) and M cone opsin (OPN1MW, *green*) antibodies indicated a normal location of rod and cone outer segments in B6 mice (A) but an ectopic location at dysplastic lesions of *Crb1^rd8^* mutants (B–D; *white* and *yellow arrowheads,* respectively). E–H. Glutamine synthetase (GLUL, *green*) antibodies revealed Müller cell soma at the center of the INL in B6 mice (E, detailed in I) but displaced toward the ONL at lesions of *Crb1^rd8^* mutants (F–H, *white arrowheads*, detailed in J– L). M–P. Purinergic receptor P2RY12 antibodies (*green*) revealed microglia at their normal location in the OPL of B6 mice but mobilized to the ONL and subretinal space among dysplastic lesions of *Crb1^rd8^* mutants (*arrowheads*). Nuclei were detected with DAPI (*blue*). Scale bars: H, 50 µm, applies to A–H and M–P; L, 25 µm, applies to I–L.

### ELM fragmentation

Pan-retinal disruption of the ELM is a distinct feature of homozygous *Crb1^rd8^* mice thought to arise from a defect in the formation and maintenance of cell-cell adhesive interactions, which depend on CRB1 [19]. This feature is observed in retinal sections of *Crb1^rd8^* mutant mice immunostained with antibodies against components of the cell adhesion machinery, such as β- actin, cadherins, PALS1, or TJP1 [19], [40], which reveal a fragmented appearance characterized by small gaps in the normally continuous ELM. It is unknown whether retinal dysplasia requires ELM fragmentation or is instead the result of an independent pathological process. To test whether fragmentation was affected by the modifier mutations, we stained retinal sections to detect TJP1. The ELM was continuous in the inferior (Fig. 7A) and superior retina (Fig. 7E) of B6 mice but exhibited gaps in the corresponding regions of B6 *rd8* (Fig. 7, B and F) and *Tvrm266* mice (Fig. 7, C and G). Strikingly, however, ELM fragmentation was rare in *Tvrm323* retinas (Fig. 7, D and H), despite an increase in the number and size of dysplastic lesions. The decrease in the number of gaps relative to B6 *rd8* was particularly noticeable in the superior retina of *Tvrm323* mice, which at one month of age had no dysplastic lesions (compare Fig. 7H and F). Note that the ELM is distorted at dysplastic lesions (for example, Fig. 7B) and therefore fragmentation cannot be evaluated at these sites. Gap counts normalized to ELM length in areas unaffected by dysplasia were determined as a measure of fragmentation (Fig. 7I, S1 Data). A statistically significant difference in this measure was observed in one or more strains compared to B6 *rd8* (one-way ANOVA, F (3, 12) = 38.90, *p* <0.0001; S1 Data). In *post hoc* analysis corrected for multiple comparisons by Dunnet’s test, statistically significant differences in fragmentation were observed in B6 and *Tvrm323* (p < 0.0001), but not *Tvrm266* (*p* = 0.091) mice, compared to B6 *rd8* mice (Fig. 7I, *asterisks*; S1 Data). These results provide further evidence that genetic modifiers affect *Crb1^rd8^* phenotypes differentially and raise the possibility that dysplasia and ELM fragmentation are independent pathological changes arising from the *Crb1^rd8^* mutation.

**Fig. 7.**
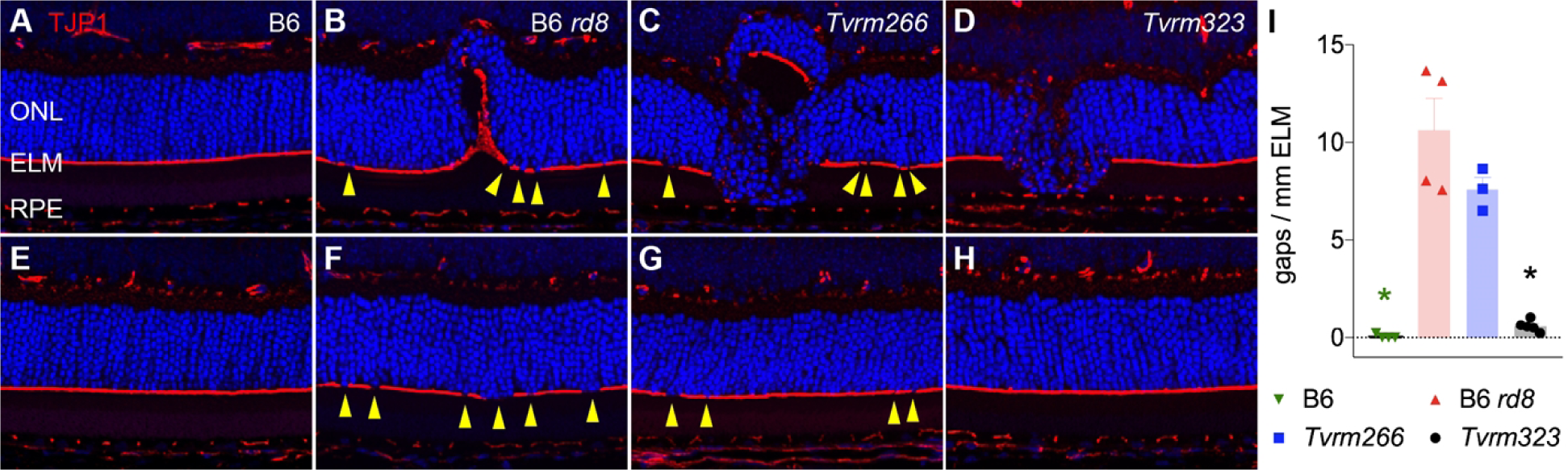
ELM fragmentation in modifier strains. Retinal sections from mice at one month of age were stained with antibodies against TJP1 (*red*) to detect the ELM and DAPI (*blue*) to detect nuclei. A–D. Inferior retina. E–H. Superior retina. A, E, B6; B, F, B6 *rd8*; C, G, *Tvrm266*; D, H, *Tvrm323.* Gaps are indicated by *yellow arrowheads.* Images are representative of n = 8 samples from each strain. H. Scale bar, 50 µm, applies to all image panels. I. Count of gaps over the full length of ELM except at dysplastic lesions, normalized to ELM length (n = 3–4). Bars indicate mean ± SEM. Asterisks, p < 0.0001, *post hoc* analysis.

### Effect of modifiers on neonatal retinal progenitor cell development

Studies of severely dysplastic *Crb1* mutant mice that also carry conditional mutations in *Crb2* targeted to photoreceptor cells, retinal progenitor cells, or Müller glia have revealed defects in early retinal development [43]. In these strains, mitotic progenitor cells of the retinal neuroblastic layer (NBL), which at P1 are normally localized within a narrow band near the RPE, are distributed ectopically throughout the NBL and even reach the ganglion cell layer [43]. To test for ectopic localization of mitotic progenitor cells in our *Crb1^rd8^* modifier strains, we examined retinal sections at P0 from B6, B6 *rd8*, *Tvrm266,* and *Tvrm323* mice stained with antibodies against phosphohistone H3, which identifies cells at mitotic prophase and metaphase (Fig. 8). Mitotic progenitor cells were similarly positioned at the NBL apical surface of all strains (Fig. 8A–D). These results indicate that mitotic progenitor cell localization at an early developmental stage is unaffected by the homozygous *Crb1^rd8^* mutation alone or in combination with the *Arhgef12* or *Prkci* modifier mutations.

**Fig. 8.**
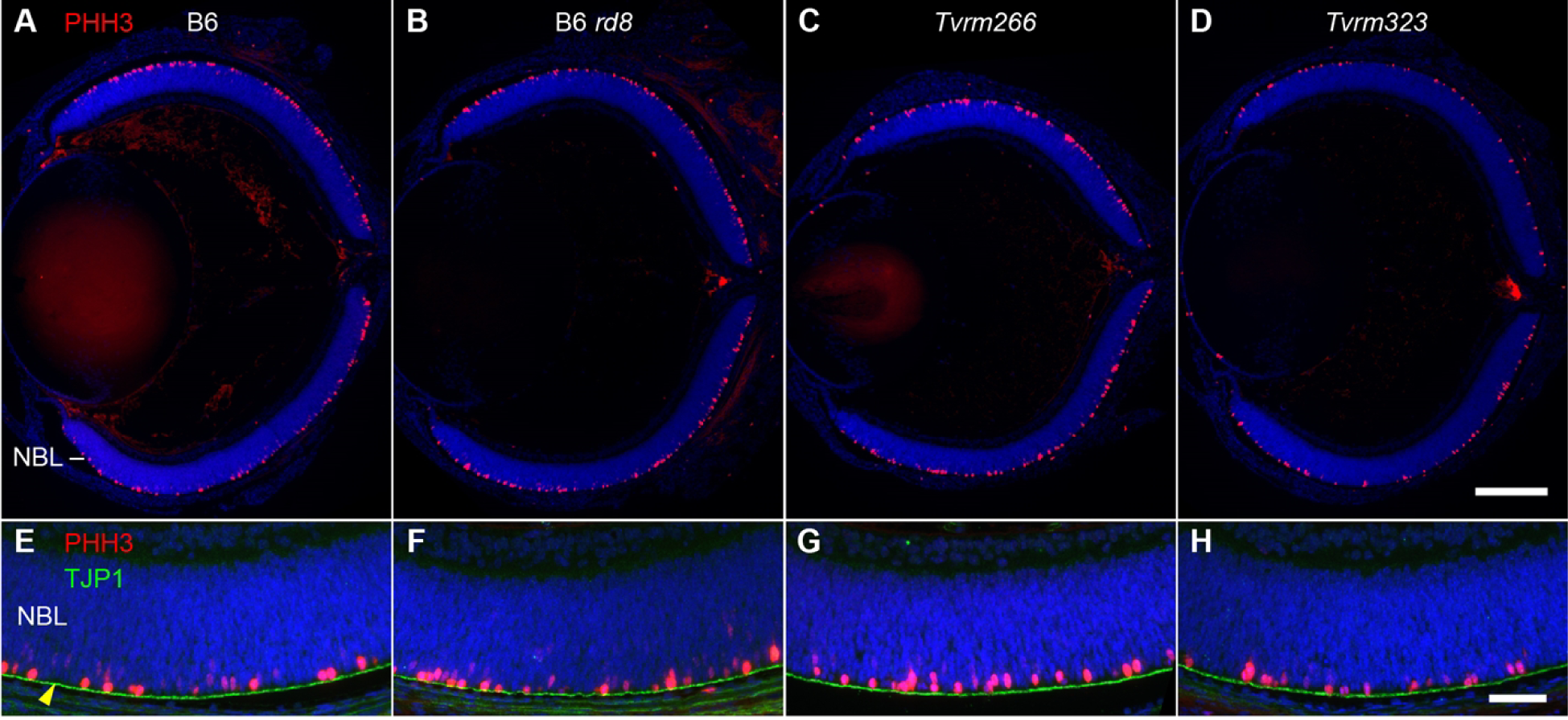
Distribution of mitotic progenitor cells at P0. A–D. Ocular sections of B6 (n = 4), B6 *rd8* (n = 5)*, Tvrm266* (n = 4), or *Tvrm323* (n = 5) mice, respectively, were probed with antibodies against phosphohistone H3 (PHH3, *red*) to detect mitotic cells and DAPI (*blue*) to detect nuclei. A similar distribution of PHH3-positive staining within the outer NBL (A, *arrow*) was observed in all strains. E–H. Higher magnification images of P0 retinas stained as in A–D and with antibody against TJP1 (*green*). The order of samples is the same as in A–D. A normal placement of progenitor cells undergoing mitosis close to the ventricular zone, delineated by TJP1 (E, *arrow*), was observed in all strains. NBL, neuroblastic layer. Scale bars: D, 0.25 mm, applies to A–D; H, 50 µm, applies to E–H.

### Loss of visual function in modifier strains

Loss of vision in *CRB1-*associated retinal disease occurs congenitally in LCA patients or progresses rapidly during early life in RP and cone-rod dystrophy patients. In comparison, mouse strains carrying a single homozygous *Crb1* mutation exhibit little to no progressive reduction in visual function as measured by electroretinography (ERG) [20], [22], [23], limiting their utility as models of *CRB1*-associated vision loss. Functional loss in mouse models is accelerated in strains combining homozygous *Crb1* mutations with modifier mutations, such as those in *Crb2* [42], [43], [52]. To assess whether a decline in visual function was accelerated in *Crb1^rd8^* modifier strains, B6, B6 *rd8*, *Tvrm266,* and *Tvrm323* mice were examined by ERG. As no significant loss of photoreceptors was noted histologically at one month of age, scotopic and photopic ERG response amplitudes as a function of flash intensity were measured at four, eight, and 12 months of age (Fig. 9A). Scotopic a- and b-wave amplitudes provide a measure of rod photoreceptor and secondary neuron function while the photopic b-wave amplitude measures secondary neuron responses to cone stimulation. Responses at each age were compared statistically to those of B6 *rd8* by mixed-effects analysis, a repeated-measures approach that allows datasets with occasionally missing values. By this analysis (S1 Data), statistically significant differences in mean scotopic a-wave amplitude were detected in one or more strains compared to B6 *rd8* mice (4 months, *p* = 0.0306; 8 months, *p* = 0.0022; 12 months; *p* = 0.0004); in mean scotopic b-wave amplitude (8 months, *p* = 0.0009; 12 months; *p* = 0.0002); and in mean photopic b-wave amplitude (8 months; *p* = 0.0034; 12 months; *p* = 0.0021). *Post hoc* analysis with Dunnet’s correction for multiple comparisons revealed a statistically significant decrease in *Tvrm266* a-waves amplitude at a single flash intensity (4 months; -1.8 log cd s/m^2^; *p* = 0.046) and in *Tvrm323* at multiple intensities (12 months; -1.4, -1.0, and-0.6 log cd s/m^2^; *p* = 0.017, 0.032, and 0.35, respectively). Parallel analysis indicated an unusual and statistically significant increase in mean *Tvrm266* b-wave amplitude (8 months; -1.4, -1.0, and-0.6 log cd s/m^2^; *p* = 0.031, 0.013, and 0.0070, respectively) as well as statistically significant decreases in mean *Tvrm323* b-wave amplitude (8 months; 1.8 log cd s/m^2^; *p* = 0.049; 12 months; -2.6, -2.2, -1.8, -1.0, and-0.6 log cd s/m^2^; *p* = .0098, 0.013, 0.029, 0.013, and 0.0098, respectively). No other comparisons, including those of B6 to B6 *rd8* were found to be statistically significant (S1 Data). These results indicate a transiently increased ERG response in *Tvrm266* mice at eight months of age, which is unexpected in light of the decreased number of photoreceptor cells predicted from histological analysis. They also indicate a loss of photoreceptor function in *Tvrm323* mice as indicated by the statistically significant decrease in scotopic a- and b-wave amplitudes at 12 months of age.

**Fig. 9.**
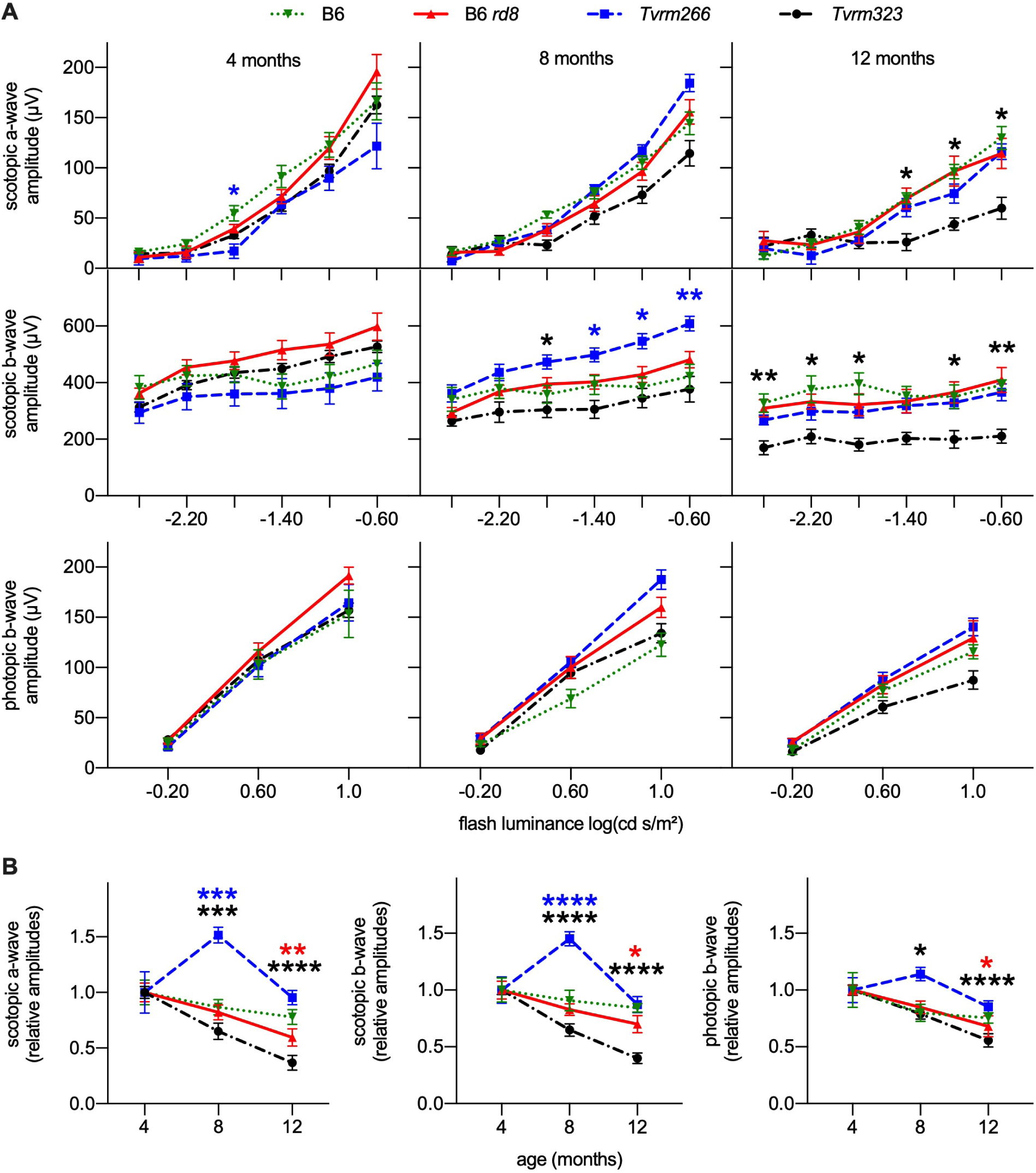
ERG analysis of B6, B6 *rd8*, *Tvrm266,* and *Tvrm323* mice at four, eight, and 12 months of age. A. Scotopic and photopic response amplitudes as a function of flash luminance. *Asterisks* show statistical significance in a comparison to to B6 *rd8* and are colored as in the legend to indicate the strain compared. * p < 0.05; ** p < 0.01. Values indicate mean ± SEM; n = 6–26. B. Response amplitudes at the highest flash luminance as a function of age, normalized to the mean amplitude at four months of age. Legend is the same as in panel A. *Asterisks* indicate statistical significance compared to the value at four months of age and are colored as in the legend in panel A. * p < 0.05, ** p < 0.01; *** p < 0.001; **** p < 0.0001.

As an alternative measure of progressive functional changes within each strain, we reanalyzed ERG response amplitudes obtained at the highest scotopic and photopic flash intensities as a function of age (Fig. 9B, S1 Data). In this analysis, amplitude values at each age were normalized and compared to the mean value at four months of age. Statistically significant differences in response amplitudes at one or more ages were detected in all strains except B6 (S1 Data; two-way ANOVA; scotopic a- and b-wave, photopic c-wave; *p* < 0.0001). *Post hoc* analysis with Dunnet’s correction for multiple comparisons revealed a statistically significant decrease in mean B6 *rd8* response amplitudes (12 months; scotopic a-wave; *p* = 0.0082; scotopic b-wave, *p* = 0.0020; photopic b-wave, *p* = 0.013); a statistically significant increase in *Tvrm266* amplitudes (8 months; scotopic a-wave; *p* = 0.0001; scotopic b-wave, p < 0.0001) and a statistically significant decrease in *Tvrm323* amplitudes (8 months; scotopic a-wave; *p* = 0.0007; scotopic b-wave, *p* < 0.0001; photopic b-wave, *p* = 0.017; 12 months; scotopic a-wave; *p* < 0.0001; scotopic b-wave, *p* < 0.0001; photopic b-wave, *p* < 0.0001). Comparison of the mean values indicated a progressive and statistically significant decrease in ERG response amplitudes in B6 *rd8* mice of 30–40% from four to 12 months of age, which was exacerbated in *Tvrm323* mice to 44–63% over the same period, consistent with degeneration of rod and cone photoreceptor cells (Fig. 9B). *Tvrm266* mice showed increased scotopic responses of 45–51% at eight months of age, a difference which was not sustained at 12 months of age. In summary, the ERG data provide evidence for functional changes in the modifier strains, which differ from each other and from B6 *rd8* mice, supporting the hypothesis that *Crb1^rd8^* modifiers differentially affect retinal function.

## Discussion

Modifier genes can alter the onset, progression, severity, and specific characteristics of monogenic ocular diseases, presenting challenges for patient diagnosis, prognosis, and treatment [7], [33]. Our finding that mutations in murine *Arhgef12* and *Prkci* modulate *Crb1^rd8^* retinal phenotypes to differing extents and with different lesion characteristics further supports the hypothesis that modifier gene variants contribute to clinical variability in *CRB1-*associated retinal dystrophy. Our results provide insight as to the pathogenic process and expand the growing network of *Crb1^rd8^* modifier genes that may ultimately lead to a greater understanding of underlying disease mechanisms that lead to the vast array of CRB1 associated disease phenotypes. The models reported here present with new retinal phenotypes, which more closely recapitulate the human disease and may be useful for pre-clinical studies.

Defects in a number of apicobasal polarity proteins have been observed to cause retinal dysplasia. Three conserved protein complexes first identified in invertebrates are thought to determine apicobasal polarity in epithelial cells and other polarized cell types: the *Drosophila* Crb-Sdt-Patj complex [mouse orthologs CRB(1–3)-MPP5- PATJ], the *Drosophila* Baz-aPKC- Par-6 complex [PARD3(or PARD3B)-PRKC(I or Z)-PARD6(A, B, or G)], and the *Drosophila* Scrib-Dlg1-L(2)gl complex [SCRIB-DLG(1–5)-LLGL(1 or 2)] [53–55]. These complexes promote the formation of tight and adherens junctions that define the apical, lateral, and basal domains of epithelial cells [56]; they also play a critical role in determining the orientation and maintenance of mitotic spindles during asymmetric cell division [57]. Importantly, members of the Rho family of small signaling GTPases control the activities of these complexes during these processes [56], [58], [59]. One family member, RHOA, is a cytoskeletal remodeling factor that plays a fundamental role in the formation and maintenance of adherens and tight junctions, which are central to apicobasal polarity [59], [72]. Another family member in *Drosophila*, Cdc42, is responsible for the membrane localization and apical accumulation of Baz, aPKC, Par- 6 and Crb, and may mediate exchange of the aPKC-Par-6 subcomplex from Baz to Crb [49]. In vertebrate models, mutation or loss of zebrafish (*Danio rerio*) orthologs of apicobasal polarity proteins EPB41L5 (zebrafish moe) [60], [61], MPP5 (nok) [62], and CRB2 (ome) [63] cause retinal developmental and lamination abnormalities similar to *Crb1-*associated retinal dysplasia. Deficits in mouse CRB1 [19], [20], [23], CRB2 [64–66], CRB1 and CRB2 combined [42], [52], [66], MPP5 [67], and PRKCI [68] have similar effects, supporting the hypothesis that apicopolarity protein defects cause retinal dysplasia.

Our results also support this hypothesis. Retinal dysplasia is more severe in *Tvrm323* mice, which express variants of two apiocobasal polarity proteins, PRKCI and CRB1, when compared to B6 *rd8* mice, in which only CRB1 is disrupted. By analogy with the apicobasal polarity processes in *Drosophila* discussed above, the PRKCI variant may alter its binding to a PARD6A, B, or G isoform and/or exchange of the resulting complex from PARD3 and CRB1. A similar, but less severe increase in retinal dysplasia is observed in *Tvrm266* mice, which are predicted to have a defect in ARHGEF12. Although ARHGEF12 is not reported to play a role in apicobasal polarity, it acts as a guanine exchange factor [69], [70] and GTPase activating protein [71] for RHOA, which influences apicobasal polarity as described above. ARHGEF12 may function similarly to ARHGEF18, a related RHOA guanine exchange factor required for apicobasal polarity in the medaka fish (*Oryzias latipes*) [73]. Strikingly, loss of ARHGEF18 in the medaka fish results in retinal dysplasia [73] similar to that caused by disruption of the CRB complex in zebrafish [60–63] and *ARHGEF18* variants cause *CRB1-*related disease in humans [74]. Thus, as in *Tvrm323,* the increased dysplasia of *Tvrm266* mice may be due to combined defects in two apicobasal polarity proteins.

Although our studies reveal genetic interaction of *Arhgef12* and *Prkci* with *Crb1,* an open question is how these interactions are mediated at the cellular level. While it is conceivable that dysplasia is due to dysfunction or loss of the encoded proteins in a single cell type, available evidence suggests that the proteins are expressed in multiple cell types. Retinal CRB1 has been detected in both Müller and photoreceptor cells [16], [19], only Müller cells [75], or separately in Müller and photoreceptor cells depending on whether CRB1-A, CRB1-B, or both isoforms were targeted by the antibody used [21]. Mouse PRKCI is widely distributed among retinal progenitor cells at the apical surface of the NBL at P0, but mainly in the inner nuclear and ganglion cell layers at P9 [68]. There is genetic evidence in mice that PRKCI functions in retinal vascular endothelial cells [76] and a search of the GEO Profiles database [77] indicated *Arhgef12* transcripts at P8 are also enriched in this cell population (GSE27238, [78]). In addition, a single-cell survey of B6 retinas at P14 [79] identified *Arhgef12* transcripts in amacrine, horizontal, and ganglion cells, astrocytes, fibroblasts, pericytes, and vascular endothelium, while *Prkci* transcripts were detected in horizontal cells (The Broad Institute Single Cell Portal, https://singlecell.broadinstitute.org/single_cell, accessed August 21, 2021). Further studies to resolve the specific cell types expressing these proteins are needed to understand the observed genetic interaction of *Arhgef12* and *Prkci* with *Crb1*.

The phenotypes observed in *Tvrm266* and *Tvrm323* mice provide important insights regarding the pathogenic process(es) leading to retinal dysplasia. First, photoreceptor cells are mislocalized and outer segment production is compromised only at dysplastic lesions. Second, in dysplastic lesions of both models and B6 *rd8* mice, Müller cell soma are displaced from their normal location in the center of the inner nuclear layer toward the RPE. Similar displacement has been observed in other murine retinal injury models [80], [81]. Third, compared to B6 control animals, microglia at lesions are mobilized toward the subretinal space in both modifier strains and in B6 *rd8* mice. Similar immune cell mobilization observed in other homozygous *Crb1^rd8^* strains [35], [38], [43] may indicate a contribution of these cells to lesion formation, although direct tests of this hypothesis have not been reported. Fourth, ELM fragmentation, which arises from a focal loss of cell-cell adhesion among Müller and photoreceptor cells, is reduced in *Tvrm323* mice compared to *Tvrm266* and B6 *rd8* mice, despite increased dysplasia. This result raises the possibility that ELM fragmentation and retinal dysplasia are independent phenotypes associated with *Crb1* mutation, in agreement with previous observations that dysplasia can be suppressed without affecting fragmentation [19] and that certain *Crb1* mutants exhibit dysplasia while retaining an ELM structure similar to that of wild-type controls [20].

Fifth, cyst-like structures within the outer nuclear layer of both models accumulate in the region of the retina where dysplastic lesions occur. These structures may be related to foveolar retinoschisis [82] or macular edema [83], often detected in *CRB1* RP patients [25], [84]. The cause of these cyst-like structures is unknown but, based on the observation of vascular leaks upon Müller cell ablation [85], they may ultimately arise from defects in junctions between Müller cells and retinal vascular beds. Finally, in *Tvrm266* and *Tvrm323* mice at both one month and one year of age, vascular structures are observed in dysplastic regions, some of which include displaced RPE cells. Vascular structures related to retinal angiomatous proliferation are observed in other *Crb1* mutant modifier strains [38], [39], [41], [43], and macular telangiectasia-like structures have been reported in a rat *Crb1* mutant [51]. Taken together, these results suggest a pathogenic process in which disruption of apicobasal polarity complexes results in abnormal activation of Müller, immune, and vascular cells, leading to edema, neovascular changes, and focal tissue remodeling observed as dysplastic lesions. Further studies of *Tvrm266* and *Tvrm323* mice may help elucidate the molecular mechanisms that underlie this process.

Our results add to a growing list of reported *Crb1^rd8^* modifier genes, which include *Cx3cr1, Mthfr, Crb2, Cygb, Jak3,* and *Nfe2l2* [35], [39–43], [45]. Analysis of modifier genes in mouse models of other diseases have been useful for assembling genetic networks that provide important clues to pathogenesis and offer new avenues for therapeutic intervention [31], [32]. As discussed above, *Crb1*, *Crb2*, *Arhgef12,* and *Prkci* may be placed in a network of apicobasal polarity genes. However, it is not obvious how the remaining modifier genes identified to date fit into this network. *Cx3cr1* encodes an immune-cell receptor for fractalkine, a signaling molecule expressed predominantly in the central nervous system and upregulated in injured neurons [86]. *Mthfr* encodes 5,10-methylenetetrahydrofolate reductase, a folate metabolic enzyme that influences serum homocysteine levels and is implicated in diverse conditions, including neurodegenerative disease [87]. *Cygb* encodes cytoglobin B, an oxygen-binding protein expressed in vascular pericytes that may protect against cellular oxidative damage [88].

*Jak3* encodes a signaling kinase that regulates myeloid and lymphoid cell development and activation [89], [90], and *Nfe2l2* encodes a transcriptional factor in many cell types that regulates oxidative stress genes [91]. Although all these genes can conceivably be associated with the phenotypes we observe, identifying additional genes that fill in missing connections will be necessary to construct a comprehensive network to explain retinal dysplasia. Our study has highlighted the utility of a sensitized mutagenesis screen to identify *Crb1^rd8^* modifier genes, which will continue to extend the network. These efforts may be amplified by the characterization of retinal dysplasia among B6N-derived strains generated by the Knockout Mouse Phenotyping Program and International Mouse Phenotyping Consortium, where every knock-out line also bears the *Crb1^rd8^* mutation [92], [93]. The potential of this second approach has been demonstrated [40].

The identification of *Arhgef12* and *Prkci* as *Crb1* modifier genes may improve disease diagnosis. Some patients possessing a single identified *CRB1* variant allele may be affected due to modifier gene variants [2]. For example, it has been suggested that *CRB2* variants may modify *CRB1* alleles in humans [42], [43]. The increased retinal dysplastic phenotype of conditional *Crb2* mutant mice carrying heterozygous *Crb1* knockout alleles [42], [43] suggests this effect could occur among patients bearing a single mutant *CRB1* allele. Identifying *Crb1* modifiers in mice may provide gene candidates for diagnostic screening of patients who retain only a single mutant *CRB1* allele. In addition, the altered disease characteristics caused by *Crb1* modifier genes, including but not limited to *Arhgef12* and *Prkci*, may aid in disease prognosis. If disruptive variants of mouse *Crb1* modifier genes are found in *CRB1* patients, a better prediction of disease onset, progression, severity and specific pathological features might be possible based on the corresponding murine phenotypes. Further, the mouse models described here exhibit new features aligned with human *CRB1* pathology, which may aid in developing therapies. The cyst-like lesions in both *Tvrm266* and *Tvrm323* mice may be useful for understanding the origin of and testing treatments for macular and foveal retinoschisis associated with *CRB1* variants [25], [82–84]. The progressive decline in rod and cone cell ERG responses in *Tvrm323* mice is faster than in mice carrying a single homozygous *Crb1* mutation, including STOCK *Crb1^rd8^* mice, and may be useful for understanding and treating functional decline in *CRB1* patients. Translational studies using these *Crb1^rd8^* modifier mouse models, which are readily available to researchers, may yield insights that improve the care of patients affected by *CRB1-*dependent retinal dystrophy.

## Methods and Materials

### Colony management and ENU mutagenesis

STOCK *Crb1^rd8^* was backcrossed to C57BL/6J (JAX stock number 000664) for seven generations to produce an incipient congenic strain, B6.Cg-*Crb1^rd8^*/Pjn (N_7_), which was bred to homozygosity. Males from this incipient congenic colony were administered weekly intraperitoneal injections of N-ethyl-N-nitrosourea (ENU) for three consecutive weeks at a concentration of 80 mg/kg per treatment [94], [95]. Following return to fertility, the mutagenized G_0_ males were backcrossed to B6.Cg-*Crb1^rd8^*/Pjn females, producing G_1_ mice which were subsequently outcrossed to unmutagenized B6.Cg-*Crb1^rd8^*/Pjn (N_7_) and resulting female G_2_ mice were backcrossed to G_1_ sires to produce G_3_ mice [46].

G_3_ mice were screened at 12 weeks of age using indirect ophthalmoscopy. Heritability of novel ocular phenotypes was established by outcrossing to B6.Cg-*Crb1*^rd8^/Pjn (N_7_) mice, producing F_1_ progeny to test for dominance. If no phenotype was observed, F_1_ mice were then intercrossed, to produce F_2_ progeny, and screened for recessive phenotypes. Once heritability was established mutant mice were assigned a number in the TVRM program [46] and lines were bred and maintained in the JAX Research Animal Facility. Mutant strains were backcrossed five generations (N_5_) to the founder strain, B6.Cg-*Crb1^rd8^/*Pjn, to remove potential unlinked mutations before samples for characterization were collected. The full designations for the modifier strains are B6.Cg-*Crb1^rd8^ Arhgef12^Tvrm266^*/Pjn and B6.Cg-*Crb1^rd8^ Prkci^Tvrm323^*/Pjn, which are abbreviated as *Tvrm266* and *Tvrm323*, respectively, and refer to a genotype that is homozygous for both *Crb1^rd8^* and the modifier allele unless otherwise indicated.

Mice were provided with NIH 6% fat chow diet and acidified water, with a 12:12 hour dark:light cycle in pressurized individual ventilation caging and were monitored regularly to maintain a pathogen-free environment. All procedures utilizing live mice were approved by the JAX Institutional Animal Care and Use Committee (protocol ACUC 99089).

### Fundus photodocumentation

Mouse pupils were dilated using 1% Cyclomydril (Alcon) prior to imaging. Isoflurane (Kent Scientific) was delivered at 2–4% with an oxygen flow rate of 0.5 liter/min to anesthetize mice during imaging. Color fundus videos (100 frames) were acquired using a Micron IV Retinal Imaging Microscope (Phoenix Technology Group), and registered, averaged, and sharpened in Fiji [96] using the ImageStabilizeMicronStack macro as described [97]. Images were further processed using a custom Fiji macro that applies the Polynomial Shading Corrector plugin (degree x = 2, degree y = 2, regularization = 50) to flatten brightness across the image and resets the range of each channel to optimize color balance.

### Gene identification

Based on observations during initial breeding, the mutations in *Tvrm266* and *Tvrm323* appeared to segregate as semidominant alleles. To determine the map position of *Tvrm323*, affected mice (enhanced spots) were outcrossed to C57BL/6NJ (B6N, JAX stock number 005304) and the resulting F_1_ progeny were backcrossed to B6N. All backcross progeny homozygous for the *Crb1^rd8^* mutation were phenotyped by indirect ophthalmoscopy. DNA from 103 backcross progeny (59 unaffected, 44 enhanced) were genotyped with 149 single nucleotide polymorphisms (B6/B6N SNP panel) spanning the genome (JAX Fine Mapping Facility). We identified genomic regions where >70% of the B6 allele co-segregated with the enhanced phenotype. Candidate genes for *Tvrm323* were identified in the minimal confidence interval on Chr 3. Gene-specific PCR amplification and subsequent Sanger sequencing confirmed a missense mutation in *Prkci.* To identify the causative mutations in *Tvrm266*, high-throughput exome sequencing was performed on a whole-exome library created from *Tvrm266* genomic DNA on a HiSeq 2000 Sequencing System (Illumina) as previously described [98]. Whole-exome sequencing of *Tvrm266* identified a nonsense mutation in *Arhgef12*, which was subsequently confirmed by Sanger sequencing. An abbreviated mapping cross, with 16 progeny (8 unaffected, 8 enhanced) confirmed the cosegregation of the affected *Tvrm266* phenotype with the *Arhgef12* mutation on Chr 9.

For strain maintenance, modifier mice were genotyped using DNA isolated from tail tips incubated with 50 mM sodium hydroxide at 95°C for about 20 min before being neutralized with 1M Tris-Cl, pH 8.0, to a final concentration of 50 mM. PCR amplification was performed in a 10 µl reaction volume containing 1X PCR Buffer (New England Biolabs), 10 µM dNTP and 0.05 U *Taq* DNA polymerase (New England Biolabs). An initial denaturation step at 97°C for 2 minutes was followed by 45 cycles of 10 seconds at 95°C, 15 seconds at 55°C, and 30 seconds at 72°C with a final 3 minute extension at 72°C. *Prkci* genotypes were determined using an allele-specific protocol combining the following primers (Eurofins Genomics): Prkci-F, TCCAGCAGTAAGTATGGGAAAC; Prkci-WT-R, GTGTGGCCATTTGCACAACA; and Prkci-MUT-R2, GTTTGGCTTGAAAAACGTGGCCATTTGCACAGTT. Depending on zygosity, this protocol yielded amplified products of 145 bp (WT), 159 bp (mutant), or both. *Arhgef12* genotypes were determined by direct sequencing. DNA samples were amplified with primers (Arhgef12-266seqF, CACACACACGTCACTGTAAA; mArhgef12-266seqR, GAGTGCCTCAATCCACATAAG). The PCR products were purified and sequenced using the reverse primer.

### Western analysis

B6, B6 *rd8, Tvrm266,* and *Tvrm323* mice (n = 4) eyes at one month of age were dissected in ice-cold PBS with proteinase inhibitor (Roche) and snap frozen in eppendorf tubes on dry ice.

Following the addition of 100 µl 1× NuPAGE^™^ LDS sample buffer/reducing agent (Invitrogen), samples were immediately homogenized using a hand-held pellet pestle motor (Kontes) on ice and briefly sonicated (Qsonica). A dilution series of the B6 samples was used to standardize the western signal. Samples were denatured at 100°C for 10 min and 5% of samples were loaded and run on a 10% Mini-Protean TGX gel (Bio-Rad). Electrophoresed proteins were transferred onto a nitrocellulose membrane using Turbo-blot Transfer system (Bio-Rad). Membranes were pre-blocked in Blotto A (5% w/v milk powder in Tris-buffered saline, 0.05% Tween [TBS-T]), and incubated with antibodies against ARHGEF12 (Santa Cruz, sc-15439), PRKCI (Novus, NBP1-84959), or GAPDH (Cell Signaling, #2118) at 4 °C overnight. Membranes were washed in TBS-T and incubated with HRP-conjugated anti-goat IgG (R&D, HAF017) or anti-rabbit IgG (Cell Signaling, 7074S) antibody in Blotto A (1:1000) for 2 h at room temperature. Following washes in TBS-T, immunoblots were visualized using the Clarity ECLWestern Blot Detection System (Bio-Rad). Protein band images in 16-bit tif format were captured using an Azure c600 bioimaging system (Azure Biosystems). The raw integrated densities of the ARHGEF12 or PRKCI bands, and the densities for GAPDH bands on the same blots, were determined using Fiji following subtraction of a constant background value evaluated between lanes. The density values for the standard series were plotted against load volume and fit to a hyperbolic curve in Prism, which was used to calculate the relative load volumes for all samples. Relative expression was determined by dividing the equivalent load volumes of the ARHGEF12 or PRKCI bands by those of the corresponding GAPDH bands. The resulting values were normalized by dividing by the mean ARHGEF12 or PRKCI values from all B6 samples.

### RNA extraction and analysis

Total RNA was extracted from B6, B6 *rd8*, *Tvrm266,* and *Tvrm323* mice (n = 5) at one month of age by homogenizing posterior eyecups in TRIzol (Thermo Fisher) using a gentleMACS dissociator (Milteny Biotec). Chloroform was then added and the resulting aqueous phase transferred to 100% ethanol. RNA clean-up was performed using an Rneasy spin column kit (Qiagen). Dnase (Qiagen) treatment was performed according to the manufacturer’s instructions. Total RNA was eluted in Rnase-free water, quantified using a NanoDrop ND1000 spectrophotometer (Thermo Fisher) and examined for quality using the TapeStation 4150 system (Agilent Technologies). cDNA was synthesized using the Superscript IV Firststrand Synthesis kit (Thermo Fisher).

To design primers for use in real time quantitative PCR, gene and coding sequences were downloaded from mouse Ensembl (https://useast.ensembl.org). Forward and reverse primers amplifying cDNA sequences of 150–250 bp and separated by large intronic regions in the genomic sequence were designed using Primer3 software (http://bioinfo.ut.ee/primer3-0.4.0/), which yielded Arhgef12 rt-F3 (AGAGCCATCAGTCACTGGACA) and Arhgef12 rt-R3 (CAGCCGTTCCTGTTCCTTC) for *Arhgef12*, and Prkci rt-F1 (GGAGTGAGGAGATGCCGAC) and Prkci rt-R1 (TCATAATATCCCCGCGGTAG) for *Prkci*.

For quantitative RT-PCR, amplification using these primers was performed with iTaq Universal SYBR Supermix (Bio-Rad Laboratories) using the CFX96 Real-time PCR Detection System (Bio-Rad). The relative fold change of gene transcripts was calculated using the comparative CT method (ΔΔC_T_), and 2^-ΔΔCT^ values were normalized to levels of *Actb*, an internal control calibrator. Melting curve analysis was evaluated to confirm accurate amplification of the target genes.

### Histological analysis

Following carbon dioxide euthanasia, eyes were enucleated, placed in chilled methanol:acetic acid:phosphate-buffered saline (PBS) (3:1:4), fixed overnight at 4°C, and stored in 70% ethanol. Fixed eyes were embedded in paraffin and 5 µm sections were stained with hematoxylin and eosin or left unstained for immunohistochemistry (see below). Eyes were oriented to yield sections parallel to the superior-inferior axis by marking the superior eye with a non-soluble paint prior to enucleation. Eyes were also oriented using the long posterior ciliary arteries, which serve as landmarks [99] and could be viewed from the back of embedded eyes illuminated through the cornea. Histosections were scanned using a NanoZoomer 2.0-HT digital slide scanner (Hamamatsu) at 40× magnification; typically 18–21 sections were captured per slide.

Sections on the slide were reviewed using NDP.view 2 software (Hamamatsu) to ensure correct orientation based on the position of the central retinal artery to one side of Bruch’s membrane opening at the optic nerve head. All sections were reviewed for the presence of pathological features, including dysplastic lesions, voids, or neovessels, and mice were graded as affected if the feature was observed in one or more sections. To assess photoreceptor cell loss from scanned images, a custom Fiji macro was first used to extract .tif images of single sections from full-resolution NanoZoomer output files, which exceded Fiji memory capabilities. A separate macro was created to draw tilted rectangular regions of interest (ROIs) encompassing a 50 µm length of retina and spaced at 0.3 mm intervals starting from the optic nerve head. Photoreceptor cell loss was determined from .tif images by manually counting photoreceptor cell nuclei within each ROI.

### Immunohistochemistry

Unstained sections of eyes were deparaffinized through a xylene/alcohol gradient series. Following deparaffinization, antigen retrieval was performed by microwave treatment for 6 min in 10 mM sodium citrate. Samples were treated with blocking solution (1:50 normal donkey serum:PBS, 0.3% v/v Triton X-100) for 30 min at room temperature and then incubated overnight at 4°C with primary antibodies against rhodopsin (MS-1233-R7, Thermo Fisher), green opsin (AB5405, EMD Millipore), glutamine synthetase (ab176562, Abcam), phosphohistone H3 (PHH3; ab14955, Abcam), and TJP1 (339100 and 61-7300, Thermo Fisher), all diluted 1:100 in blocking solution. Slides were washed using 1× PBS prior to application of secondary antibodies at 1:200 in blocking solution for two h at room temperature. Nuclei were stained using DAPI (10 µg/ml in PBS) for 5 min. Negative controls, in which primary antibodies were omitted during the incubation stage, were performed in parallel. All samples were mounted in Vectashield (Vector Laboratories), coverslipped and imaged using a Zeiss Axio Observer.z1 fluorescence microscope (Carl Zeiss Microscopy) with an ApoTome2 attachment. To count ELM gaps, ApoTome2 stacks encompassing the full thickness of the stained section and covering the entire retina were acquired and processed to yield a maximum intensity projection using Zen 2.6 software (Carl Zeiss). The resulting images were viewed in Fiji and gaps counted manually. The ELM was manually fit to a spline curve, avoiding dysplastic regions identified in the DAPI channel. The gap count was divided by the total ELM length calculated from the fitted curves. To assess the distribution of mitotic nuclei at P0, images were acquired with a 5× objective using a Leica DMLB fluorescence microscope and analyzed in Fiji with a macro that compares the intensity of PHH3 fluorescence near the apical boundary of the NBL to that in the full NBL [100].

### ERG analysis

ERG analysis was carried out using a Espion V6 ColorDome Multifocal system (Diagnosys) essentially as described [101]. Mice at four, eight, and 12 months of age were dark-adapted overnight and pupils were dilated using1% atropine sulfate (Akorn) or 1% cyclopentolate (Akorn). Mice were anesthetized using ketamine (Covetrus) and xylazine (Akorn) diluted in normal saline and administered by intraperitoneal injection at a final dose of 80 and 16 mg/kg of body weight, respectively. Anesthetized mice were placed on the heated platform of the Espion system. Gold loop electrodes were placed on the corneal surface of each eye along with Goniovisc hypromellose ophthalmic solution (HUB Pharmaceuticals), Gonak (Akorn), or Refresh (Allergan). Small needle electrodes beneath the skin at the top of the head and at the tail served as reference and system ground, respectively. Eyes of dark-adapted mice were presented with a six-step protocol of increasing light intensity measuring mainly rod-dependent responses, followed by light adaptation and a three-step protocol measuring cone-dependent responses. The flash luminance (cd s m^-2^), number of sweeps, and time between sweeps (s) for the dark-adapted series was 0.0025, 5, 5; 0.006, 5, 5; 0.016, 5, 5; 0.04, 5, 10; 0.1, 5, 15; and 0.25, 5, 15, respectively. Following a 10-min light adaptation at 110 cd m^-2^, the corresponding parameters for the light-adapted series were 0.63, 10, 1; 4, 10, 1; and 10, 20, 1 on a background illumination of 110 cd m^-2^. Response amplitudes were determined from the average of multiple sweeps as follows: scotopic a-wave, pre-flash baseline to the negative trough; scotopic b-wave, negative trough to positive peak within 40–120 ms following the flash; photopic b-wave, negative trough to positive peak within 40–60 ms following the flash. The mean response amplitude from both eyes of each mouse was used for statistical analysis; if data were missing from one eye, only one eye was used.

### Statistical analysis

Genotype-phenotype associations to identify the causative mutation and to determine the association of strain with pathological features were assessed with Fisher’s Exact test using JMP, Version 15 or 16 (SAS Institute). Quantitative immunoblotting and qRT-PCR data were analyzed by one-way ANOVA and Student’s t-test, respectively, using Prism (GraphPad).

Genotype-phenotype association in epistasis crosses were analyzed by linear regression modeling. Heterozygous and homozygous genotypes were independently coded to assess allelic dominance in both main and interaction effects. Significance was estimated by the ratio of F-statistics between interactive and additive models, with *p*-values derived from the Fisher-Snedecor distribution. ERG results were analyzed in Prism using a mixed effect model to accommodate missing data and to allow the use of repeated measures; further analysis of normalized data at the highest ERG flash intensities was performed in Prism using two-way ANOVA. A significance threshold of *p* < 0.05 was used for all experiments.

## Acknowledgements

We thank Melissa Berry for nomenclature review. We also thank JAX Scientific Research Services, including the Computational Sciences, Genome Technologies, Genetic Engineering Technologies, Histopathology, and Microscopy Services.

